# Fulgor: A fast and compact *k*-mer index for large-scale matching and color queries

**DOI:** 10.1101/2023.05.09.539895

**Authors:** Jason Fan, Noor Pratap Singh, Jamshed Khan, Giulio Ermanno Pibiri, Rob Patro

## Abstract

The problem of sequence identification or matching — determining the subset of references from a given collection that are likely to contain a query nucleotide sequence — is relevant for many important tasks in Computational Biology, such as metagenomics and pan-genome analysis. Due to the complex nature of such analyses and the large scale of the reference collections a resource-efficient solution to this problem is of utmost importance. The reference collection should therefore be pre-processed into an *index* for fast queries. This poses the threefold challenge of designing an index that is efficient to query, has light memory usage, and scales well to large collections.

To solve this problem, we describe how recent advancements in associative, order-preserving, *k*-mer dictionaries can be combined with a compressed inverted index to implement a fast and compact *colored de Bruijn* graph data structure. This index takes full advantage of the fact that unitigs in the colored de Bruijn graph are *monochromatic* (all *k*-mers in a unitig have the same set of references of origin, or “color”), leveraging the *order-preserving* property of its dictionary. In fact, *k*-mers are kept in unitig order by the dictionary, thereby allowing for the encoding of the map from *k*-mers to their inverted lists in as little as 1 + *o*(1) bits per unitig. Hence, one inverted list per unitig is stored in the index with almost no space/time overhead. By combining this property with simple but effective compression methods for inverted lists, the index achieves very small space.

We implement these methods in a tool called `Fulgor`. Compared to `Themisto`, the prior state of the art, `Fulgor` indexes a heterogeneous collection of 30,691 bacterial genomes in 3.8× less space, a collection of 150,000 *Salmonella enterica* genomes in approximately 2 × less space, is at least twice as fast for color queries, and is 2 − 6× faster to construct.

**2012 ACM Subject Classification:** Applied computing → Bioinformatics

## 1 Introduction

At the core of many metagenomic and pan-genomic analyses is *read-mapping*, the atomic operation that assigns observed sequence reads to putative genome(s) of origin. A wide range of methods have been developed for mapping reads to large collections of reference genomes. Of note, alignment-based methods, though accurate [19, 22], are relatively computationally intensive as they must provide the ability to *locate* the read on each genome. A queried read must, with low edit-distance, be matched with a sub-string of some reference genome in the collection. For alignment, the index is also required to report the position of this match. As a matter of fact, alignment against hundreds or even tens of thousands of reference genomes can be impractically slow and simply require too much space in practice.

Fortunately, *alignment-free* techniques have become popular and widespread for metagenomic analyses [42, 26, 41, 34, 39, 35]. These methods generally work by avoiding alignment altogether, and replacing it with strategies for matching (exactly or approximately) sub-strings, signatures, or sketches between the queries and the referenced sequences. Ideally, good matching heuristics can assign or match a query against the correct reference with high precision while also retaining high recall (i.e., being sensitive to sequencing error or small divergence between the query and the reference). One particular type of alignment-free method for assigning reads to compatible references that has recently gained substantial traction is *pseudoalignment* [6, 38, 36, 20]. While tremendous progress has been made in supporting alignment-free methods for metagenomic analyses, continued development of ever more efficient indexing methods is required for such analyses to scale to tens, even hundreds, of thousands of bacterial reference genomes.

A practical data structure that is suitable for alignment-free matching methods is the *colored de Bruijn graph*, a graph where each node corresponds to a *k*-mer in a reference collection and is annotated with a *color*, the set of references in which it occurs. Bifrost [15] and Metagraph [17] are two efficient approaches that index the colored de Bruijn graph and support the *k*-mer-to-color query. Recently, Alanko et al. [2] developed `Themisto`, an index for alignment-free matching (and specifically pseudoalignment) that substantially outperforms these prior methods in the context of indexing and mapping against large collections of genomes. Compared to Bifrost, `Themisto` uses practically the same space, but is faster to build and query. Compared to the fastest variant of Metagraph, `Themisto` offers similar query performance, but is much more space-efficient; on the other hand, `Themisto` is much faster to query than Metagraph-BRWT, the most-space efficient variant of Metagraph.

### 1.1 Contributions

We describe how recent advancements in associative, order-preserving, *k*-mer dictionaries [29, 28] can be combined with a compressed inverted index to implement a fast index over the *colored compacted* de Bruijn graph (ccdBG). Leveraging the *order-preserving* property of its dictionary, our index takes full advantage of the fact that unitigs in this variant of the ccdBG are *monochromatic* — i.e., all *k*-mers in a unitig have the same set of references of origin, or “colors”. In fact, *k*-mers are kept in unitig order, and our index takes advantage of the ability of our associative dictionary to store the unitigs in any order. Reordering the unitigs so that all color-equivalent unitigs are adjacent in the index allows the construction of a map from *k*-mers to their corresponding colors that uses only 1 + *o*(1) bits per unitig. Our index combines this property with a simple but effective hybrid compression scheme for inverted lists (colors) to require little space. By storing unitigs and keeping *k*-mers in unitig order, our index also supports very fast streaming queries for consecutive *k*-mers in a read, and additionally allows efficient implementation of skipping heuristics that have previously been suggested to speed up pseudoalignment [6]. We implemented our index in a C++17 tool called `Fulgor`, which is available at https://github.com/jermp/fulgor.

Compared to `Themisto` [2], the prior state of the art, `Fulgor` indexes a heterogeneous collection of 30,691 bacterial genomes in 3.8× less space, a collection of 150,000 *Salmonella enterica* genomes in approximately 2× less space, is at least twice as fast at query time, and even 2 − 6 × faster to construct.

Perhaps unsurprisingly, the rapid development of novel indexing data structures has been accompanied by novel and custom strategies for matching and assigning reads to colors (i.e., reference sets) and algorithms that each make different design choices and trade-offs. Many of these strategies can be considered as a form of pseudoalignment. Having been iterated on since its introduction [6], the term “pseudoalignment” has come to describe a family of efficient heuristics for read-to-color assignment, rather than a single concept or algorithm. Prior methods have taken either *exhaustive* approaches that queries every *k*-mer on a read (previously termed *exact* pseudoalignment [20, 2]) or have implemented *skipping* based approaches that skip the query of “redundant” consecutive *k*-mers that likely map to the same set of reference genomes [6, 13]. To our knowledge, the precise details of the types of skipping heuristics used in the latter methods — including those adopted by the initial pseudoalignment method — have been discussed only in passing. Complete details, instead exist only in the source code of the corresponding tools. To shed light on these algorithms, we provide a more structured discussion of how these algorithms are designed. Using `Fulgor`, we implement two previously proposed variants and benchmark them.

## 2 Preliminaries

In this section, we first formalize the problem under study here. We then describe a modular indexing layout that solves the problem using the interplay between two well-defined data structures. Lastly we describe the properties induced by the problem and how these are elegantly captured by the notion of *colored compacted de Bruijn graph*.

### 2.1 Problem definition

▸ **Problem 1** (Colored *k*-mer indexing problem). *Let ℛ* = {*R*_1_, …, *R*_*N*_} *be a collection of references. Each reference R*_*i*_ *is a string over the DNA alphabet* Σ ={ *A, C, G, T* }. *We want to build a data structure (referred to as the* index*) that allows us to retrieve the set* `Color` (*x*) = { *i* | *x* ∈ *R*_*i*_ } *as efficiently as possible for any k-mer x* ∈ Σ^*k*^. *Note that* `Color` (*x*) = ∅ *if x does not occur in any reference*.

Hence, we call the set `Color` (*x*) the *color* of the *k*-mer *x*.

### 2.2 Modular indexing layout

In principle, Problem 1 could be solved using an old but elegant data structure: the *inverted index* [44, 33]. The inverted index, say ℒ, stores explicitly the ordered set `Color` (*x*) for each *k*-mer *x* ∈ ℛ. What we want is to implement the map *x* → `Color` (*x*) as efficiently as possible in terms of both memory usage and query time. To this end, all the distinct *k*-mers of ℛ are stored in an *associative* dictionary data structure, 𝒟. Suppose the dictionary 𝒟 stores *n k*-mers. To implement the map *x* → `Color` (*x*), the operation that 𝒟 is required to support is `Lookup` (*x*) which returns ⊥ if *k*-mer *x* is not found in the dictionary or a unique integer identifier in [*n*] = {1, …, *n*} if *x* is found. Problem 1 can then be solved using these two data structures — 𝒟 and ℒ — thanks to the interplay between `Lookup` (*x*) and `Color` (*x*): logically, the index stores the sets {`Color` (*x*) }_*x*∈ℛ_ in compressed format in the order given by `Lookup` (*x*).

To our knowledge, all prior solutions proposed in the literature that fall under the “color-aggregative” classification [21], are incarnations of this *modular indexing framework* and, as such, require an efficient *k*-mer dictionary joint with a compressed inverted index. For example, `Themisto` [2] makes use of the *spectral* BWT (or `SBWT`) data structure [1] for its *k*-mer dictionary, whereas Metagraph [17] implements a general scheme to compress metadata associated to *k*-mers which is, in essence, an inverted index.

### 2.3 The colored compacted de Bruijn graph and its properties

Problem 1 has some specific properties that one would like to exploit to implement as efficiently as possible the modular indexing framework described in Section 2.2. First, consecutive *k*-mers share (*k*− 1)-length overlaps; second, co-occurring *k*-mers have the same color. A useful, standard, formalism that describes these properties is the *colored compacted de Bruijn graph* (abbreviated “ccdBG”).

Given the collection of references ℛ, the (node-centric) de Bruijn graph (dBG) of ℛ is a directed graph whose nodes are all the distinct *k*-mers of ℛ and there is an edge connecting node *u* to node *v* if the (*k*− 1)-length suffix of *u* is equal to the (*k*− 1)-length prefix of *v*. We refer to *k*-mers and nodes in a (node-centric) dBG interchangeably; likewise, a path in a dBG spells the string obtained by “glueing” together all the *k*-mers along the path. Thus, unary (i.e., non-branching) paths in the graph can be collapsed into single nodes spelling strings that are referred to as *unitigs*. The dBG arising from this compaction step is called the compacted dBG (cdBG). Lastly, the *colored* compacted dBG is obtained by logically annotating each *k*-mer *x* with its color, `Color` (*x*), and only collapsing non-branching paths with nodes having the same color.

Below, we notate *n* to be the number of distinct *k*-mers of ℛ and *m* to be the number of unitigs {*u*_1_, …, *u*_*m*_ }of the ccdBG induced by the *k*-mers of ℛ. The unitigs of the ccdBG that we consider have the following key properties.

1. *Unitigs are contiguous subsequences that spell references in ℛ*. Each distinct *k*-mer of ℛ appears once, as sub-string of some unitig of the cdBG. By construction, each reference *R*_*i*_∈ ℛ can be a *tiling* of the unitigs — a sequence of unitig occurrences that spell out *R*_*i*_ [11]. Joining together *k*-mers into unitigs reduces their storage requirements. In Sections 3.1 and 3.2, we show how this property can be exploited to make indexes compact. In Section 4, we show how this property can be exploited to make queries fast.
2. *Unitigs are monochromatic*. The *k*-mers belonging to the same unitig *u*_*i*_ all have the same color. Thus, we shall use `Color` (*u*_*i*_) to denote the color of each *k*-mer *x* ∈*u*_*i*_. We note that this property holds only if one considers *k*-mers appearing at the start or end of reference sequences to be *sentinel k*-mers that must terminate their containing unitig [23, 18], and that such conventions are not always adopted [15, 8].
3. *Unitigs co-occur and share colors*. Unitigs often have the same color (i.e., occur in the same set of references) because they derive from conserved sequences in indexed references that are longer than the unitigs themselves. We indicate with *M* the number of distinct color sets 𝒞 = {*C*_1_, …, *C*_*M*_ }. Note that *M* ≤*m* and that in practice there are dramatically more unitigs than there are distinct colors. We use `ColorID` (*u*_*i*_) = *j* to indicate that unitig *u*_*i*_ has color *C*_*j*_. As a consequence, each *k*-mer *x* ∈ *u*_*i*_ has color *C*_*j*_.

In this work our goal is to design an index that takes full advantage of these key properties.

## 3 Index description

In this section we describe a modular index that implements a colored compacted de Bruijn graph (ccdBG) and fully exploits its properties described in Section 2.3. We adopt the modular indexing framework from Section 2.2 — comprising a *k*-mer dictionary 𝒟 and an inverted index ℒ — to work seamlessly over the *unitigs* of the ccdBG. We extend the ideas from Fan et al. [11] for the modular indexing of *k*-mer positions to *k*-mer colors.

Our strategy is to first map *k*-mers to unitigs using a dictionary 𝒟, and then map unitigs to their colors 𝒞 = {*C*_1_, …, *C*_*M*_ }. By *composing* these mappings, we obtain an efficient map directly from *k*-mers to their associated colors. The colors themselves in 𝒞 are stored in compressed form in a inverted index ℒ. Figure 1 offers a pictorial overview of how we orchestrate these different components in the index. The goal of this section is to describe how these mapping steps can be performed efficiently and in small space.

**Figure 1.**
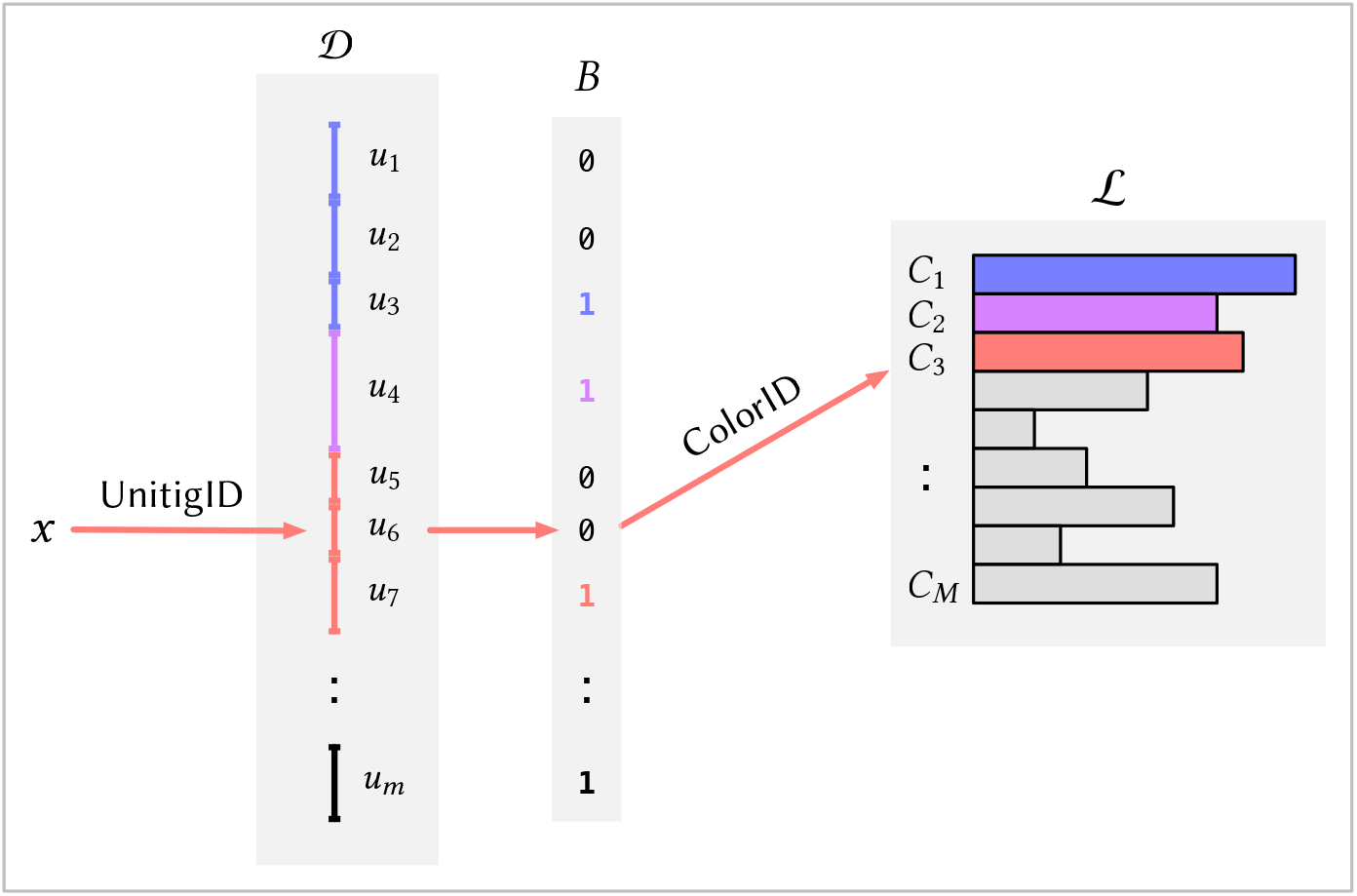
A schematic picture of the index described in Section 3, highlighting the interplay between the *k*-mer dictionary 𝒟, the bit-vector *B*, and the inverted index ℒ. The red arrows show how the index is queried for a *k*-mer *x*, assuming that *x* occur in unitig *u*_6_ and has color *C*_3_. The *k*-mer *x* is first mapped by 𝒟 to its unitig *u*_6_ via the query `UnitigID`(*x*) = 6. Then we compute `ColorID`(*u*_6_) = `Rank`_1_ (6, *B*) + 1 = 2 + 1 = 3 and lastly retrieve *C*_3_ from ℒ.

### 3.1 The *k*-mer dictionary: mapping *k*-mers to unitigs

For a *k*-mer dictionary, we use the `SSHash` data structure [29, 28], which fulfills the requirement described in Section 2.2, in that it implements the query `Lookup` (*x*) for any *k*-mer *x* efficiently and in compact space. This is achieved by storing the unitigs explicitly (i.e., as contiguous, 2-bit encoded strings) in some prescribed order so that a *k*-mer *x* occurring in some unitig *u*_*i*_ can be quickly located using a minimal perfect hash function [31] built for the set of the *minimizers* [37] of the *k*-mers. Laying out unitigs in this principled manner also enables very efficient streaming query. That is, when querying consecutive *k*-mers from input reads, the query for a given *k*-mer can often be answered very efficiently given the query result from its predecessor, since it often shares the same minimizer and frequently even occupies the very next position on the same unitig as its predecessor. We refer the interested reader to [29, 28] for a complete overview of `SSHash`.

Even more importantly for our purposes, a query into the `SSHash` dictionary returns, among other quantities, `UnitigID` (*x*) = *i*, the ID of the unitig containing the *k*-mer *x*, as a byproduct of `Lookup` (*x*). For any *k*-mer occurring in ℒ, `UnitigID` (*x*) = *i* is an integer in [1..*m*]. This map from *k*-mers to unitigs will be exploited in the subsequent sections.

### 3.2 Mapping unitigs to colors

Now that we have an efficient map from *k*-mers to unitigs, i.e., the operation `UnitigID` (*x*), we must subsequently map unitigs to distinct colors. That is, we have to describe how to implement the operation `ColorID` (*u*_*i*_) for each unitig *u*_*i*_. Since each `ColorID` (*u*_*i*_) is an integer in [1..*M*], we could implement `ColorID` (*u*_*i*_) just by storing `ColorID` (*u*_1_), …, `ColorID` (*u*_*m*_) explicitly in an array of ⌈log_2_(*M*) ⌉-bit integers. We show how to do this in just 1 + *o*(1) bits per unitig rather than ⌈ log_2_(*M*) ⌉ bits per unitig.

We do so by exploiting another key property of `SSHash` : the unitigs it stores internally can be permuted in any desired order without impacting the correctness or efficiency of the dictionary. This was already noted and exploited in [28] to compress *k*-mer abundances. Similarly, here we sort the unitigs by `ColorID` (*u*_*i*_), so that all the unitigs having the same color are stored consecutively in `SSHash`. To compute `ColorID` (*u*_*i*_), all that is now required is a `Rank` _1_ query over a bit-vector *B*[1..*m*] where *B*[*i*] = 1 if `ColorID` (*u*_*i*_) ≠ `ColorID` (*u*_*i*+1_) and *B*[*i*] = 0 otherwise, 1 ≤*i < m*. Assuming the last bit of *B* is always set, then it follows that *B* has exactly *M* bits set. The operation `Rank` _1_(*i, B*) returns the number of ones in *B*[1, *i*) and can be implemented in *O*(1) time, requiring only *o*(*m*) additional bits as overhead on top of the bit-vector [40, 30]. This means that `ColorID` (*u*_*i*_) can be computed in *O*(1) as `Rank` _1_(*i, B*) + 1.

We illustrate this unitig to color ID mapping in Figure 1. In this toy example, `ColorID` (*u*_6_) = 3 can be computed with `Rank` _1_(6, *B*) + 1 = 2 + 1 because there are two bits set in *B*[1, 6) — each marking where previous groups of unitigs with the same color end. Therefore, according to *B*, unitigs {*u*_1_, *u*_2_, *u*_3_} all have the same color as also {*u*_5_, *u*_6_, *u*_7_} ; *u*_4_’s color is not shared by any other unitig instead.

### 3.3 Compressing the colors

The inverted index ℒ is a collection of sorted integer sequences {*C*_1_, …, *C*_*M*_ }, whose integers are drawn from a universe of size *N* (the total number of references in the collection ℛ). There is a plethora of different methods that may be used to compress integer sequences (see, e.g., the survey [33]). Testing the many different techniques available on genomic data is surely an interesting benchmark study to carry out. Here, however, we choose to adopt a simple strategy based on the widespread observation that effective compression appears to require using different strategies based on the density of the sequence *C*_*i*_ to be compressed (ratio between |*C*_*i*_| and *N*).

We therefore implement the following *hybrid* compression scheme:

1. For a sparse color set *C*_*i*_ where |*C*_*i*_| */N <* 0.25, we adopt a delta-gap encoding: the differences between consecutive integers are computed and represented via the universal Elias’ δ code [10].
2. For a dense color set *C*_*i*_ where | *C*_*i*_ | */N >* 0.75, we first take the complementary set of *C*_*i*_, that is, the set 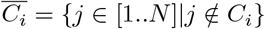, and then compress 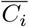 as explained in 1. above.
3. Finally, for a color set *C*_*i*_, that does not fall into either above density categories, we store a characteristic bit-vector encoding of *C*_*i*_ — a bit-vector *b*[1..*N*] such that *b*[*j*] = 1 if *j* ∈ *C*_*i*_ and *b*[*j*] = 0 otherwise.

The compressed representations of all sequences are then concatenated into a single bit-vector, say *sequences*. An additional sorted sequence, *offsets*[1..*M*], is used to record where each sequence begins in the bit-vector *sequences*, so that the compressed representation of the *i*-th sequence begins at the bit-position *offsets*[*i*] in *sequences*, 1≤ *i* ≤*M*. The *offsets* sequence is compressed using the Elias-Fano encoding [9, 12] and takes only a (very) small part of the whole space of ℒ unless the sequences are very short.

This hybrid encoding scheme is similar in spirit to the one also used in `Themisto` which, in turn, draws inspiration from Roaring bitmaps [7]. However, our choice of switching to the complementary set when |*C*_*i*_| approaches *N* turns out to be a very effective strategy, especially for pan-genome data, where a striking fraction of integers in ℒ are indeed covered by these extremely dense sets (see also Table 4 from Section 5).

### 3.4 Construction

`Fulgor` is constructed by directly processing the output of `GGCAT` [8], an efficient algorithm to build ccdBGs using external memory and multiple threads. Importantly, `GGCAT` provides the ability to iterate over unitigs grouped by color. Therefore, `Fulgor` construction just requires a single scan of the unitigs in the order given by `GGCAT`. `SSHash` is built on the set of unitigs, each distinct color is compressed as described in Section 3.3, and the bit-vector *B* is also built during the scan.

## 4 Pseudoalignment algorithms

The term *pseudoalignment*, originally coined by Bray et al. [6] and developed in the context of RNA-seq quantification, has been used to describe many different algorithms and approaches, several of which do not actually comport with the original definition. Specifically, Bray et al. [6] define a “pseudoalignment of a read to a set of transcripts, *T*” as “a subset, *S* ⊆*T*, without specific coordinates mapping each base in the read to specific positions in each of the transcripts in *S*”. The goal of such an approach then becomes to determine, for a given read, the *set* of indexed reference sequences with which the read is *compatible*, where, in the most basic scenario, the compatibility relation can be determined entirely by the presence/absence of *k*-mers in the read in specific references.

Given any index of *k*-mer colors, a variety of different pseudoalignment algorithms can be implemented that rapidly map given reads to compatible reference sequences according to a set of heuristics. Below, we describe four pseudoalignment algorithms that fall into two categories: (1) *exhaustive* methods that retrieves the color of every *k*-mer on a given read (as described in [2]), and (2) *skipping* heuristics that skip or jump over *k*-mers during pseudoalignment that are likely to be *uninformative* (i.e., to have the same color as the *k*-mer that was just queried).

### 4.1 Exhaustive methods

For a given query sequence *Q*, exhaustive approaches return colors with respect to a set of *k*-mers of *Q, K*(*Q*), that map to a non-empty color (i.e., each *k*-mer *x* ∈ *K*(*Q*) if found in the dictionary 𝒟).

#### Full-intersection

The first of the two exhaustive approaches, the *full-intersection* method, simply returns the intersection between all the colors of the *k*-mers in *K*(*Q*). Algorithm 1 in the Appendix (page 19) shows how this query mode is implemented in `Fulgor`. In the current implementation, `Fulgor` has a generic intersection algorithm that can work over *any* compressed color sets, provided that an iterator over each color supports two primitives — `Next` and `NextGEQ` (*x*), respectively returning the integer immediately after the one currently pointed to by the iterator and the smallest integer which larger-than or equal-to *x*. (We point the reader to [25] and [33] for details.)

#### Threshold-union

The second algorithm, which we term the *threshold-union* approach, relaxes the full-intersection method to trade off precision for increased recall. Instead of requiring a reference to be compatible with *all* mapped *k*-mers, the threshold-union method requires a reference to be compatible with a user defined proportion of *k*-mers. Given a parameter *τ* ∈ (0, 1], this method returns the set of references that occur in *at least s · τ* returned (i.e., non-empty) *k*-mer colors, where *s* can be either chosen to be *s* = |*K*(*Q*) | (the number of positive *k*-mers only) or *s* = |*Q*| − *k* + 1 (the total number of *k*-mers in *Q*). `Themisto` [2] implements the variant with *s* = |*K*(*Q*) | (called the “hybrid” method), whereas both Bifrost [15] and Metagraph [17] use *s* = |*Q*| *k* + 1. In fact, the latter approach of simply looking up all of the *k*-mers in a query, and requiring a specified fraction of them to match, is a long-standing strategy that predates the notion of pseudoalignment [43, 42]. In the following, we assume *s* = |*K*(*Q*) | is used by the threshold-union algorithm, unless otherwise specified. The pseudocode for this query mode is given in Algorithm 3 in the Appendix (page 20).

In practice, both the aforementioned exhaustive methods are efficient to compute for two reasons. First, intersections, thresholding, and unions are easy to compute because colors are encoded as monotonically increasing lists of reference IDs. Second, for `Fulgor` in particular, querying *every k*-mer for its color can be performed in a highly-optimized way via *streaming* queries to `SSHash`. In the streaming setting, `SSHash` may skip comparatively slow hashing and minimizer lookup operations because it stores *unitig* sequences contiguously in memory. When sequentially querying adjacent *k*-mers on a read that are also likely adjacent on indexed unitigs, it can rapidly lookup and check *k*-mers that are cached and adjacent in memory (we refer the reader to [29] for more details).

### 4.2 *Skipping* heuristics

For even faster read mapping, pseudoalignment algorithms can implement heuristic *skipping* approaches that avoid exhaustively querying all *k*-mers on a given read. These skipping heuristics make the assumption that whenever a *k*-mer on a read is found to belong to a unitig, subsequent *k*-mers will likely map to the same unitig and can therefore be skipped, since they will be uninformative with respect to the final color assigned to the query (i.e., the intersection of the colors of the mapped *k*-mers).

Bray et al. [6] first described such an approach, where a successful search that returns a unitig *u* triggers a skip that moves the search position forward to either the end of the query or the implied distance to the end of *u* (whichever is less). Subsequent searches follow the same approach as new unitigs are discovered and traversed in the query. Later, other tools extended or modified the proposed skipping heuristics, and introduced “structural constraints”, which take into account the co-linearity and spacing between matched seeds on the query and on the references to which they map [13]. In contrast to `Themisto`, `Fulgor` indexes unitigs and has rapid access to the topology of the ccdBG. `Fulgor` thus permits efficient implementation of pseudoalignment algorithms with skipping heuristics since, due to the underlying capabilities provided by `SSHash`, it can rapidly find *k*-mers bookending unitig substrings because `SSHash` can explicitly map *k*-mers to their offsets (positions) in indexed unitig sequences.

In general, pseudoalignment methods that implement skipping heuristics must specify what steps the algorithm will take in *all* scenarios, not just what should happen when search proceeds as expected. In practice, implementations for resolution strategies are complicated and difficult to describe succinctly in prose, and prior work has only discussed these important details in passing. Here, using the depicted scenarios in Figure 2, we provide a more structured (though certainly not exhaustive) discussion of possible design choices that can be made. These design choices impact the performance of the pseudoalignment algorithm, both in terms of how many *k*-mers it queries (and, hence, its speed), and in how many distinct color sets it collects (and, hence, the actual compatibility assignment it makes).

**Figure 2.**
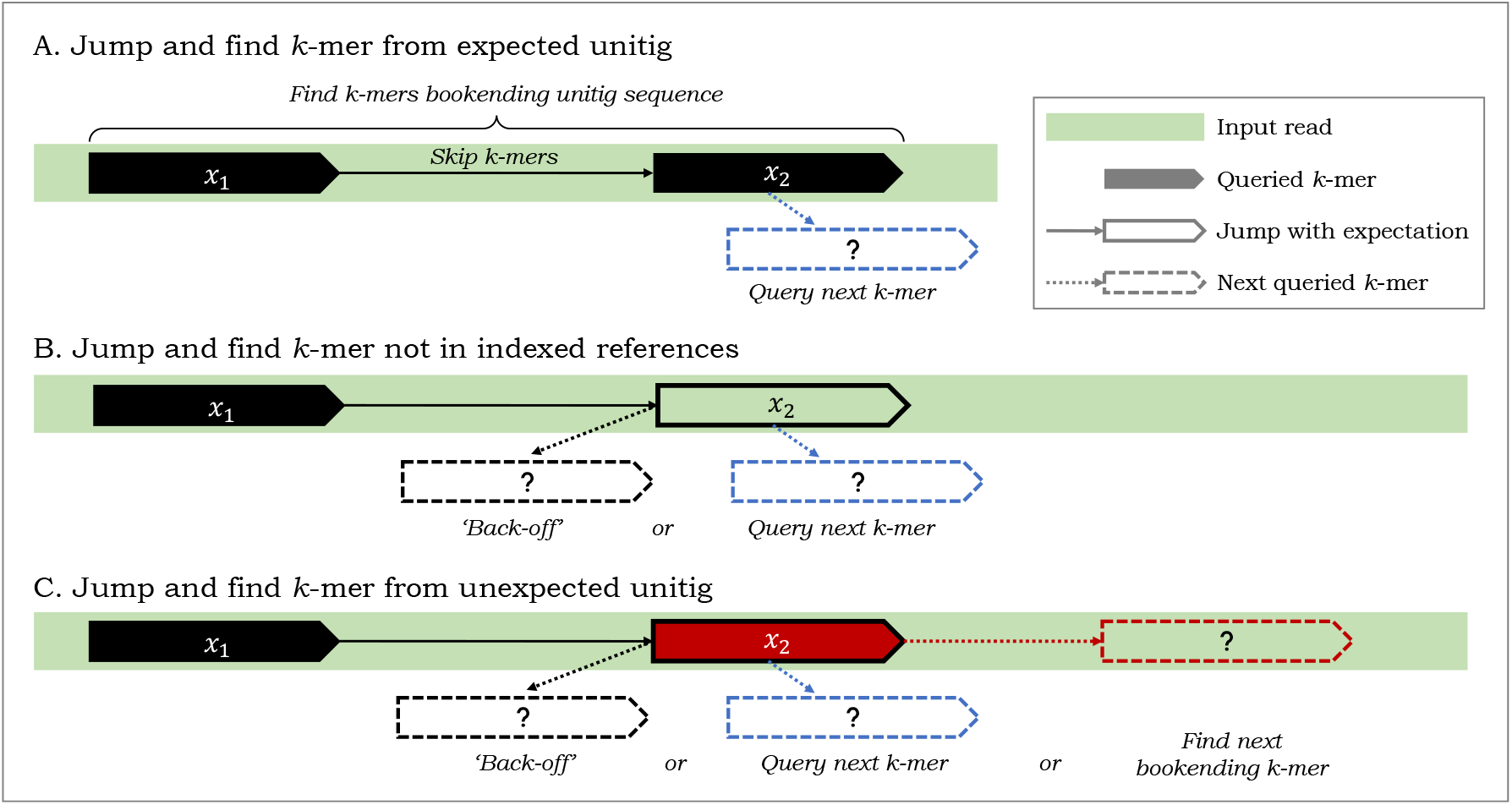
Some relevant design choices for pseudoalignment with skipping heuristics that *jump* and skip *k*-mers on a given read. After *k*-mer *x*_1_ is queried and found to map to a “black” unitig, an algorithm can jump to query the *k*-mer *x*_2_ on input read, where the number of *k*-mers skipped is given by the length of the black unitig. (A) In the ideal scenario, *x*_2_ maps to the black unitig sequence and *k*-mers *x*_1_ and *x*_2_ are found to bookend this unitig sequence as it appears on the read. (B) If *x*_2_ misses the index, an algorithm can *back-off* to an earlier *k*-mer on the read to find a *k*-mer bookending a shorter subsequence of the black unitig; or it may just query the next *k*-mer. (C) If *x*_2_ maps to a different “red” unitig, an algorithm has an alternative, aggressive, heuristic option to jump and find the next *k*-mer bookending the red unitig sequence.

#### Jump and find *k*-mer in expected unitig

Before the first matching *k*-mer of a read is found, there is relatively little difference between exhaustive and heuristic pseudoalignment approaches; subsequent *k*-mers are queried until the read is exhausted or some *k*-mer is found in the index. At this point, however, heuristic skipping methods diverge from the exhaustive approaches. At a high level, when a *k*-mer on a read is found to map to a unitig, skipping heuristics make an assumption that said unitig appears wholly on the read. A pseudoalignment algorithm then jumps, on the read, to what would be the last *k*-mer on the unitig sequence occurring on the given read (i.e., a bookending *k*-mer). Scenario A in Figure 2 depicts when this assumption is correctly made. Moving left-to-right on a given read, if a *k*-mer on the *left* is found to occur on the unitig depicted in black color in the figure (referred to as the “black” unitig henceforth), an algorithm can then skip a distance given by the length of the black unitig and jump to a *k*-mer to the right that also maps to the black unitig and bookends it. Doing so, an algorithm can assume that all *k*-mers bookended by these two queried *k*-mers map to the black unitig, avoid querying *k*-mers in-between, and instead continue query the next *k*-mer on the read (indicated in dashed lines in blue).

#### Jump and miss *k*-mer

In practice however, the implemented skipping heuristics are not so simple. This is because, when skipping *k*-mers according to unitig lengths, the resulting *k*-mer that an algorithm jumps to may not necessarily map to the unitig it *expects*. In scenario B, an algorithm jumps to a *k*-mer on a read, expecting it to map to a black unitig, but finds that it does not correspond to any indexed *k*-mer. Here, an algorithm can make several choices, and in fact, current skipping heuristics make two distinct choices in this scenario. It can ignore this missed *k*-mer and simply query the next *k*-mer after the position that was jumped to (in blue). Or, it can take a more conservative approach and implement a *back-off* scheme to look for another *k*-mer that maps to the black unitig. An algorithm can back-off and jump a lesser distance, and such a back-off approach can happen once or can be recursive or iterative until some termination condition is satisfied.

#### Jump and find *k*-mer in *un*-expected unitig

In scenario C, an algorithm that jumps to a *k*-mer but finds that it maps to a *different* (red) unitig than expected. Here, we suggest three choices an algorithm can make. Like in scenario B, an algorithm can back-off to find another *k*-mer mapping to the black unitig or it can query the next *k*-mer after the jumped position. Alternatively, it can take a new more aggressive approach and jump to a *k*-mer on the read where it expects to find the end of an occurrence of the red unitig.

In this work, we have retrofit the pseudoalignment with skipping algorithms from Kallisto [6]^1^ and Alevin-fry [13] ^2^ to make use of `Fulgor`, rather than the distinct indexes atop which they were implemented in their original work. Using `Fulgor`, we compare their resulting pseudoalignments, along with those from the full-intersection and threshold-union approaches, in a simple simulated scenario in Section 5.4.

## 5 Results

In this section, we report experimental results to assess `Fulgor`’s construction time/space, index size, and query speed. Throughout the section, we compare `Fulgor` to `Themisto` [2], which has been shown to outperform other methods that build similarly capable indexes (namely Bifrost [15] and Metagraph [17]) in terms of speed and space. We build `Themisto` indexes using the fastest configuration, i.e., without sampling of *k*-mer colors in the `SBWT` (build option −d1), as done by the authors in [2]. Not sampling *k*-mer colors yields slightly larger indexes but makes `Themisto` faster to query. For our largest benchmarked reference collection (150,000 genomes), potential space savings from sampling is not significant anyway because the space required to store distinct colors dominate the overall space. We also use `Themisto`’s default color set representation (i.e., without Roaring bitmaps). For both `Fulgor` and `Themisto`, we set the *k*-mer size to *k* = 31.

### Datasets

We follow the experimental methodology of [2] and build `Fulgor` over subsets of *Salmonella enterica* genomes (up to 150,000 genomes) from [5], to demonstrate `Fulgor`’s effectiveness when indexing collections of similar reference sequences. We also consider a heterogeneous collection of 30,691 genomes of bacterial species representative of the human gut [14] (as also benchmarked in our previous work [11]). We report some summary statistics for the indexed ccdBGs in Table 1.

**Table 1.**
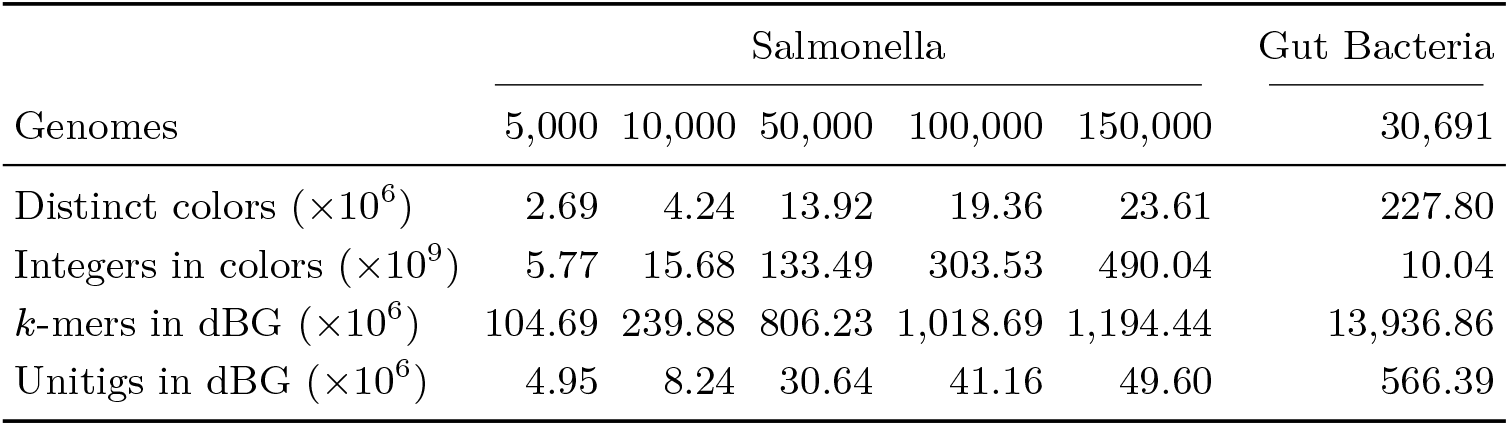
Summary statistics for the tested collections. The row “Integers in colors” reports the total number of reference IDs that are required to encode all colors — i.e., the sum set sizes for all colors, ∑_*i*_|*C*_*i*_|.

### Hardware and software

All experiments were run on a machine equipped with Intel Xeon Platinum 8276L CPUs (clocked at 2.20GHz), 500 GB of RAM running Ubuntu 18.04.6 LTS (GNU/Linux 4.15.0). `Fulgor` is available at https://github.com/jermp/fulgor. For the experiments reported here we use v1.0.0 of the software, compiled with gcc 11.1.0. For `Themisto`, we use the shipped compiled binaries (v3.1.1).

### 5.1 Construction time and space

Construction time and peak RAM usage is reported in Table 2 for the different datasets evaluated. Both tools use `GGCAT` to build the ccdBG. However, `Fulgor` is 2− 6× faster, and typically consumes much less memory during construction. This is because `Themisto` spends most of its time and memory building the color mapping. However, the analogous component of `Fulgor` is just a bit vector, demarcating groups of unitigs with the same color, that is built via a linear scan of the unitigs produced by `GGCAT`.

**Table 2.** Total index construction time and GB of memory (max. RSS), as reported by /usr/ bin/time with option -v. The reported time includes the time taken by GGCAT to build the ccdBG (using 48 processing threads) and the time to serialize the index on disk.

|  | Genomes | Fulgor |  | Themisto |  |
| --- | --- | --- | --- | --- | --- |
|  |  | hh:mm | GB | hh:mm | GB |
| Salmonella | 5,000 | 00:04 | 12.91 | 00:11 | 12.97 |
|  | 10,000 | 00:09 | 23.60 | 00:25 | 23.58 |
|  | 50,000 | 01:13 | 43.76 | 02:32 | 96.00 |
|  | 100,000 | 02:56 | 73.54 | 06:25 | 202.42 |
|  | 150,000 | 04:36 | 136.94 | 10:00 | 323.10 |
| Gut Bacteria | 30,691 | 02:27 | 115.05 | 15:35 | 327.72 |

Figure 3 shows, instead, `Fulgor`’s construction time breakdown for some illustrative datasets. We distinguish between three phases in the construction: (1) running `GGCAT`, (2) compressing the colors and, (3) building `SSHash`. While `GGCAT` and color compression take most of the construction time on the Salmonella pangenomes, building `SSHash` is the most expensive step on the Gut Bacteria collection. This is consistent with the statistics reported in Table 1. Here, there are far more integers to compress in the Salmonella collections whereas the Gut Bacteria collection contains one order of magnitude more *k*-mers. This suggests that one could achieve even faster construction for `Fulgor` if the colors are compressed in parallel with the `SSHash` construction (currently, these two phases are sequential).

**Figure 3.**
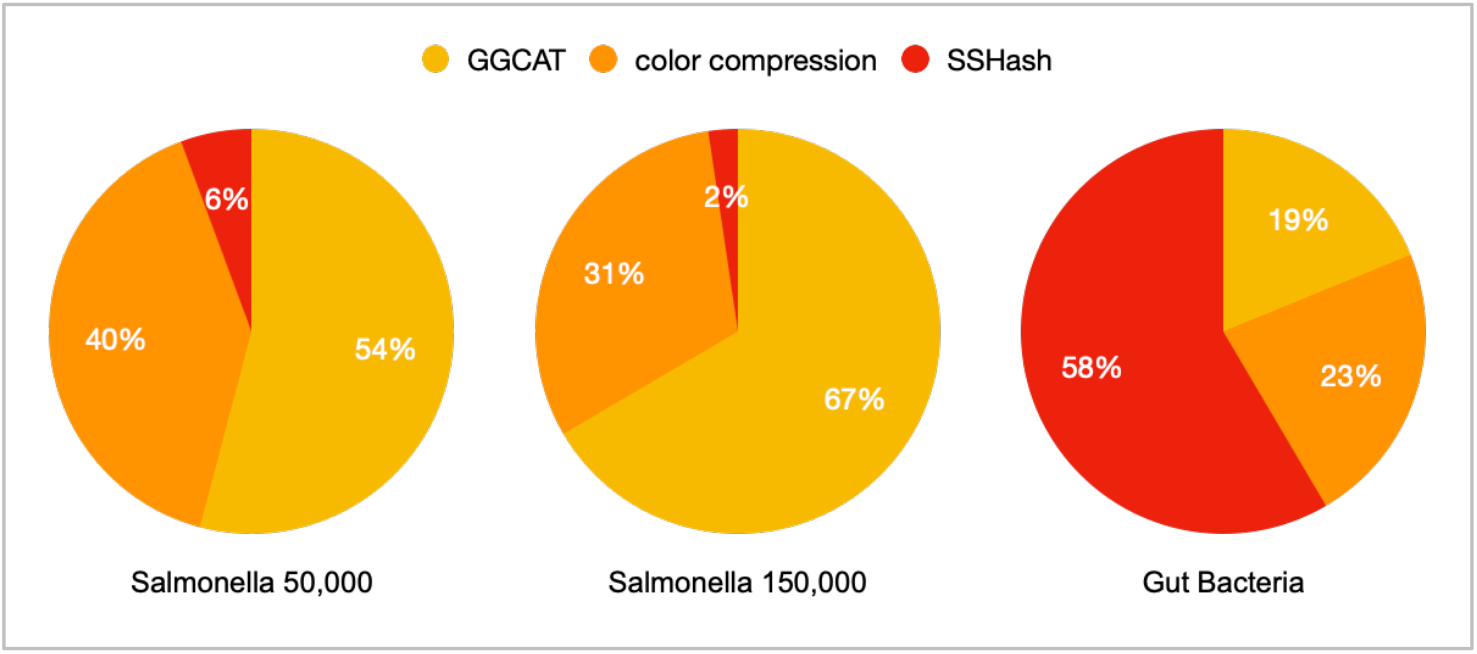
Construction time breakdown for Fulgor.

### 5.2 Index size

When indexing collections of Salmonella genomes, `Fulgor` is consistently ≈2× smaller than `Themisto` as apparent from Table 3. For example, on largest collection comprising 150,000 genomes, `Fulgor` takes 70.66GB whereas `Themisto` takes 133.63GB. This remarkable space improvement is primarily due to the more effective color compression scheme adopted by `Fulgor`. This leads to, for example, 48% less space to encode colors for the 150,000 collection of Salmonella genomes. Looking at Table 4, we highlight that for all indexed Salmonella reference collections, approximately 50% of all encoded integers in the distinct colors belong to colors that are *at least* 90% dense. For such extremely dense colors, the complementary encoding strategy described in Section 3.3 is very effective: only ≈ 0.2 bpi are required to encode them in all benchmarked indexes. In fact, even for our largest collection of 150,000 Salmonella genomes, encoding *all* integers in *all* colors requires only 1.120 bpi.

**Table 3.** Index space in GB, broken down by space required for indexing the *k*-mers in a dBG (SSHash for Fulgor, and the SBWT for Themisto); and data structures required encode colors and map *k*-mers to colors.

|  | Genomes | Fulgor |  |  | Themisto |  |  |
| --- | --- | --- | --- | --- | --- | --- | --- |
|  |  | dBG | Colors | Total | dBG | Colors | Total |
| Salmonella | 5,000 | 0.16 | 0.59 | 0.75 | 0.14 | 1.82 | 1.96 |
|  | 10,000 | 0.35 | 1.66 | 2.01 | 0.32 | 4.78 | 5.09 |
|  | 50,000 | 1.26 | 17.03 | 18.30 | 1.07 | 36.89 | 37.96 |
|  | 100,000 | 1.72 | 40.70 | 42.44 | 1.35 | 81.82 | 83.17 |
|  | 150,000 | 2.03 | 68.60 | 70.66 | 1.58 | 132.05 | 133.63 |
| Gut Bacteria | 30,691 | 21.23 | 15.45 | 36.77 | 18.33 | 121.08 | 139.41 |

**Table 4.** Average bits/int (bpi) spent for representing colors whose density is (*i*× 10)% of *N*, for *i* = 1, …, 10. The first two columns for each collection, “lists” and “ints”, report the percentage of lists (i.e., colors) and integers (stored reference identifiers) that belong to all colors within a given density. The last row, “Total bpi”, is comprehensive of the space spent for the integer lists themselves and the space spent for the offsets delimiting the lists’ boundaries.

| Density | Salmonella |  |  |  |  |  |  |  |  |  |  |  |  |  |  | Gut Bacteria |  |  |
| --- | --- | --- | --- | --- | --- | --- | --- | --- | --- | --- | --- | --- | --- | --- | --- | --- | --- | --- |
| | $N = 5,000$ | | | $N = 10,000$ | | | $N = 50,000$ | | | $N = 100,000$ | | | $N = 150,000$ | | | $N = 30,691$ | | |
|  | lists | ints | bpi | lists | ints | bpi | lists | ints | bpi | lists | ints | bpi | lists | ints | bpi | lists | ints | bpi |
| 0–10% | 38.88 | 2.00 | 4.66 | 46.15 | 1.81 | 4.64 | 70.96 | 2.62 | 6.00 | 76.52 | 3.14 | 6.16 | 79.23 | 3.27 | 6.32 | 99.99 | 99.99 | 12.05 |
| 10–20% | 6.04 | 2.11 | 2.66 | 4.83 | 1.93 | 2.66 | 3.74 | 2.84 | 3.05 | 2.82 | 2.61 | 3.77 | 2.54 | 2.68 | 3.92 | 0.00 | 0.00 | 0.00 |
| 20–30% | 4.70 | 2.69 | 2.86 | 4.44 | 2.93 | 2.81 | 2.69 | 3.50 | 3.24 | 2.32 | 3.66 | 3.41 | 2.09 | 3.76 | 3.46 | 0.00 | 0.00 | 0.00 |
| 30–40% | 3.13 | 2.55 | 2.88 | 4.27 | 4.02 | 2.87 | 1.90 | 3.43 | 2.89 | 1.57 | 3.49 | 2.88 | 1.40 | 3.51 | 2.88 | 0.00 | 0.00 | 0.00 |
| 40–50% | 4.05 | 4.25 | 2.23 | 3.32 | 4.04 | 2.22 | 1.81 | 4.25 | 2.22 | 1.44 | 4.14 | 2.23 | 1.29 | 4.19 | 2.23 | 0.00 | 0.00 | 0.00 |
| 50–60% | 4.13 | 5.30 | 1.83 | 3.54 | 5.29 | 1.81 | 1.82 | 5.24 | 1.82 | 1.42 | 4.99 | 1.82 | 1.24 | 4.94 | 1.82 | 0.00 | 0.00 | 0.00 |
| 60–70% | 3.98 | 6.07 | 1.54 | 4.24 | 7.44 | 1.54 | 2.04 | 6.94 | 1.53 | 1.59 | 6.61 | 1.54 | 1.40 | 6.59 | 1.54 | 0.00 | 0.00 | 0.00 |
| 70–80% | 5.53 | 9.72 | 0.94 | 4.86 | 9.91 | 0.93 | 2.33 | 9.13 | 1.08 | 1.87 | 8.96 | 1.14 | 1.64 | 8.91 | 1.15 | 0.00 | 0.00 | 0.00 |
| 80–90% | 5.80 | 11.52 | 0.47 | 3.71 | 8.57 | 0.47 | 3.03 | 13.43 | 0.56 | 2.49 | 13.65 | 0.63 | 2.09 | 12.95 | 0.66 | 0.00 | 0.00 | 0.00 |
| 90–100% | 23.77 | 53.80 | 0.15 | 20.65 | 54.07 | 0.14 | 9.67 | 48.63 | 0.19 | 7.94 | 48.76 | 0.21 | 7.07 | 49.21 | 0.21 | 0.00 | 0.00 | 0.00 |
| Total bpi | 0.817 |  |  | 0.848 |  |  | 1.020 |  |  | 1.072 |  |  | 1.120 |  |  | 12.32 |  |  |

Unsurprisingly, `Fulgor` also uses less space than `Themisto` to support the `ColorID` operation. We recall from Section 3.2 that `Fulgor` requires only 1 + *o*(1) bits per unitig by design. This amounts to a negligible space usage compared to the overall index size. For example, while `Themisto` requires 7.26GB to map *k*-mers to color IDs for 150,000 Salmonella genomes, our strategy just takes 7.75 MB.

When indexing a *heterogeneous* collection, e.g., the 30,691 bacterial genomes [14], with many more unique *k*-mers, the space advantage `Fulgor` has over `Themisto` is even more apparent. First, the overall size of `Fulgor` is 3.8 × smaller (36.77GB versus 139.41GB). Second, `Fulgor`’s near optimal approach of mapping *unitigs* to colors instead of *k*-mers to colors is dramatically more efficient, requiring only 88MB compared to `Themisto`’s 91GB. `Themisto`, by using the `SBWT`, organizes *k*-mers based on their *colexicographical* order and requires ⌈log_2_(*M*) ⌉ bits per sampled *k*-mer to record the color IDs. Here, the `SBWT` must record colors for each of the 13.9 billion distinct *k*-mers *and* their reverse complement. In contrast, `Fulgor` uses `SSHash` that maintains *k*-mers in unitig order and requires only 1 + *o*(1) bits per unitig to map *all k*-mers from the same unitig to a single color. Although not the default behavior, `Themisto` can optionally sample *k*-mer colors to avoid storing one color ID per each *k*-mer. Clearly, the sampling scheme reduces space usage at the expense of some overhead at query time by requiring an implicit walk in the dBG. While this sampling strategy can be quite effective when the underlying *k*-mer set displays long unitigs, allowing the sampling of the terminal *k*-mers of a non-branching path [2], it is unlikely to be similarly effective in a highly-branching and fragmented graph like the one underlying this heterogeneous dataset. On the contrary, our index does not have this issue *by design* and can thus scale to more heterogenous collections using small space.

### 5.3 Query speed

To compare query speed, we benchmark `Fulgor` and `Themisto` using both low- and high-hit rate read-sets, i.e., read-sets for which we have a low and high number of positive *k*-mers respectively. Precisely, we use the files containing the first read of the following paired-end libraries: SRR896663^3^ with 5.7 ×10^6^ reads, SRR801268^4^ with 6.6× 10^6^ reads, and ERR321482^5^ with 6.8 ×10^6^ reads.

In Table 5 we report the result of the comparison using the full-intersection method (Algorithm 1). We repeated the same experiment using the threshold-union method (Algorithm 3) with parameter *τ* = 0.8 as this is the preferred query mode in `Themisto`. However, we did not observe any appreciable difference compared to the full-intersection method in terms of query speed.

**Table 5.** Total query time as elapsed time reported by `/usr/bin/time`, using 16 processing threads for both indexes. The read-mapping output is written to `/dev/null` for this experiment. We also report the mapping rate in percentage (fraction of mapped read over the total number of queried reads). Results are relative to the full-intersection query mode. All reported timings are relative to a second run of the experiment, when the index is loaded faster from the disk cache. For each workload, we indicate the run accession number.

| (a) low-hit, Salmonella, SRR896663 |  |  |  | (b) high-hit, Salmonella, SRR801268 |  |  |  |
| --- | --- | --- | --- | --- | --- | --- | --- |
| Genomes | Mapping rate | Fulgor | Themisto | Genomes | Mapping rate | Fulgor | Themisto |
|  |  | mm:ss | mm:ss |  |  | mm:ss | mm:ss |
| 5,000 | 1.27 | 00:09 | 00:32 | 5,000 | 89.53 | 01:16 | 03:50 |
| 10,000 | 13.86 | 00:10 | 00:36 | 10,000 | 89.76 | 02:26 | 07:35 |
| 50,000 | 32.61 | 00:25 | 01:05 | 50,000 | 91.31 | 19:15 | 41:25 |
| 100,000 | 34.09 | 00:45 | 01:39 | 100,000 | 91.52 | 35:50 | 82:14 |
| 150,000 | 34.01 | 01:06 | 05:02 | 150,000 | 91.61 | 42:30 | 120:08 |

| (c) low-hit, Gut Bacteria, SRR896663 |  |  |  | (d) high-hit, Gut Bacteria, ERR321482 |  |  |  |
| --- | --- | --- | --- | --- | --- | --- | --- |
| Genomes | Mapping rate | Fulgor | Themisto | Genomes | Mapping rate | Fulgor | Themisto |
|  |  | mm:ss | mm:ss |  |  | mm:ss | mm:ss |
| 30,691 | 11.90 | 0:57 | 2:58 | 30,691 | 92.98 | 01:16 | 02:45 |

In a low-hit rate workload where a small proportion of reads map to the indexed references, `Fulgor` is much faster than `Themisto`. In this scenario, we expect many queried *k*-mers to not occur in the indexed references. When *k*-mers are absent from the index, no color needs to be retrieved and only the *k*-mer dictionary is queried. Here, `Fulgor` is faster than `Themisto` because its reliance on the fast streaming query capabilities of `SSHash`. It is worth noting here that in any *streaming* setting where consecutive *k*-mers are queried, `Fulgor` can fully exploit the monochromatic property of unitigs in ways which `Themisto` cannot. Queries to `SSHash` have very good locality compared to the `SBWT` because adjacent *k*-mers in unitigs are stored contiguously in memory. Further, streaming queries to `SSHash` can be very efficiently cached and optimized. When looking up consecutive *k*-mers, `SSHash` can skip slow hashing operations and instead perform fast comparisons of *k*-mers stored in cached positions pointing to adjacent addresses in memory.

In a high-hit rate workload, `Fulgor` also outperforms `Themisto`, but by a smaller margin, since most of the time is now spent in performing the intersection between colors. It is interesting to note that the workloads can be processed significantly faster (by both tools) on the Gut Bacteria collection: this is a direct consequence of the fact that the lists being intersected are much shorter on average for the Gut Bacteria compared to the Salmonella collections. This is evident from Table 4: essentially all lists are just 10% dense, i.e., have length at most ⌈30, 691*/*10⌉ *<* 3,070.

We also note that part of the slowdown seen for `Themisto` is due to the time spent in loading the index from disk to RAM, which takes at least twice as `Fulgor`’s because of its larger index size.

### 5.4 Comparison of pseudoalignment algorithms on simulated data

To analyze the accuracy of the underlying pseudoalignment algorithms, we perform additional testing with read sets simulated using the `Mason` [16] simulator. To analyze how mapping and hit rates affect query speed, we simulate a varying proportion of “positive” reads from indexed reference sequences and generate “negative” reads from the human chromosome 19 from the CHM13 v2.0 human genome assembly [24]. We use `Fulgor` to compare the four mapping algorithms described in Section 4.

From Table 6, we see that at various proportions of ground truth positive reads (simulated reads deriving from indexed references), all mapping methods have a true positive rate (TPR), i.e., total reads correctly mapped over the total ground truth positives, greater than 95%. This high sensitivity for all four methods is to be expected since all methods simply check for *k*-mer’s membership to references of origin and do not consider *k*-mer positions in references. One main drawback of eliding positions, heuristically avoiding “locate” queries, and entirely ignoring *k*-mers that are not present in the index, is also clear. All methods incur approximately a 30% false positive rate (FPR), i.e., total reads spuriously mapped over the total ground truth negatives. As is expected, the threshold-union method incurs a slightly higher FPR compared to other methods (30% compared to 27% for other methods) because of its less strict criteria only requiring references to be compatible with *τ* fraction of mapped *k*-mers instead of *all k*-mers.

**Table 6.** Quality of pseudoalignment algorithms querying 100,000 simulated reads against 50,000 Salmonella genomes indexed with `Fulgor`. We vary the percentage of *positive* reads simulated from indexed Salmonella genomes by diluting queried read sets with *negative* reads simulated from a reference human transcriptome. We consider a mapped positive read (deriving from indexed references) to be a *true positive* if the reference of origin is in the returned set of compatible references; and a mapped negative read (deriving from human chromosome 19) to be a *false positive*. We denote true and false positive rates (%) to be TPR and FPR, respectively. For the threshold-union method, we use *τ* = 0.8.

| % Positive | Full-intersection |  | Threshold-union |  | Kallisto |  | Alevin-fry |  |
| --- | --- | --- | --- | --- | --- | --- | --- | --- |
|  | TPR | FPR | TPR | FPR | TPR | FPR | TPR | FPR |
| 90% | 95.0 | 27.0 | 97.7 | 30.0 | 95.0 | 27.0 | 95.1 | 27.0 |
| 70% | 95.1 | 27.0 | 97.7 | 30.0 | 95.1 | 27.0 | 95.1 | 27.0 |
| 25% | 95.1 | 27.0 | 97.7 | 30.0 | 95.2 | 27.0 | 95.2 | 27.0 |
| 10% | 95.5 | 27.0 | 97.8 | 30.0 | 95.5 | 27.0 | 95.5 | 27.0 |

In these benchmarks, we find very little difference in terms of TPR and FPR between the exhaustive methods and skipping heuristics. These results also gesture at one desirable and one undesirable quality of these methods. First, skipping heuristics correctly assume and successfully skip *k*-mers that likely occur on the same unitig and have the same color. Likewise, they have the *potential* to be even more sensitive than the full-intersection method, as they do not, in general, search for every *k*-mer in a query, and can thus avoid scenarios where variation or sequencing errors in a query cause spurious matches to the index, shrinking or eliminating the set of references appearing in the final color assigned to the query. In fact, in a small-scale test, Alanko et al. [2] report that Kallisto’s skipping heuristic results in a small but persistent increase of approximately 0.03% in the mapping rate. However, all four of the pseudoalignment methods evaluated here suffer from a high FPR and low precision. Better algorithms to lower FPR and improve precision without lowering sensitivity too much should be investigated in future work. Such improvements may be possible by adding back information about the *reference positions* where *k*-mers from the query match, incorporating structural constraints [13] or other such restrictions atop the color intersection rule. Yet, those approaches are more computationally involved, require the index to support locate queries, and also substantially diverge from “pseudoalignment” as traditionally understood. Regardless, we highlight here that `Fulgor` more easily enables implementing skipping and unitig-based heuristics compared to other methods that do not explicitly store unitig sequences and keep *k*-mers in unitig order. In fact, `Fulgor` implicitly maintains additional information regarding the *local structural consistency* of *k*-mers. For example, with `Fulgor`, one can easily check if consecutive *k*-mers are valid on an indexed unitig or check if consecutive unitigs on a read have valid overlaps, in an attempt to reduce the FPR.

## 6 Conclusions and future work

We introduce `Fulgor`, a fast and compact index for the *k*-mers of a colored compacted de Bruijn graph (ccdBG). Using, `SSHash`, an order-preserving *k*-mer dictionary, `Fulgor` fully exploits the monochromatic property of unitigs in ccdBGs. `Fulgor` implements a very succinct map from unitigs to colors, taking only 1 + *o*(1) bits per unitig. Further, `Fulgor` applies an effective hybrid compression scheme to represent the set of distinct colors. Across all benchmarked scenarios, `Fulgor` outperforms `Themisto`, the prior state-of-the-art in terms of space *and* speed. There is still room for improvement in future work. We discuss some promising directions below.

In terms of speed, we remark that when processing a high-hit workload, the overall runtime is dominated by the time required to *intersect* the colors. As explained in Section 4.1, `Fulgor` currently implements a generic intersection algorithm that only requires two primitive operations, namely `Next` and `NextGEQ` (see also Appendix A). But this is not the only paradigm available for efficient intersection. We could, for example, try approaches that exploit different indexing paradigms, such as Roaring [7] and Slicing [27], that are explicitly designed for fast intersections. These alternative approaches may be significantly faster especially on the high-hit workloads.

Another possible optimization is to implement a caching scheme for frequently occurring and/or recently intersected colors. Caching the uncompressed or intersection-optimized versions of frequently occurring color sets, or previously computed intersections, could speed up query processing substantially when many reads map to the same set of colors.

In terms of space, one property that `Fulgor` does not yet exploit is the fact that *many* unitigs in the ccdBG share *similar colors* — i.e., co-occur in many reference sequences. This is so because unitigs arising from conserved genomic sequences will share similar occurrence patterns. In a related line of research, [3] developed a method that efficiently compresses distinct, but highly-correlated colors, through a variant of referential encoding. Specifically, they compute a minimum spanning tree (MST) on a subgraph of the color graph induced by the ccdBG, and encode a color by recording its differences with respect to its parent in the MST. This vastly reduces the space required to encode the color set when many similar colors exist, as we would expect in a pangenome, and fast query speed can be retained through color caching. Another related approach would be to resort to clustering similar colors and encoding all colors within a cluster with respect to a cluster representative color [32]. Likewise, although not specifically designed to compress colors, Metagraph and its variants can exploit similarity between colors using a general compression scheme that records differences in stored metadata (in this case, the colors) between adjacent *k*-mers [17]. We note that, since the colored *k*-mer indexing problem is *modular* (Section 2.2), novel relational compression techniques for the set of distinct colors can be developed and optimized independently of the other components of the index.

Finally, in our experiments with simulated data analyzing the quality of pseudoalignment algorithms from Section 5.4, we find higher than desirable false positive rates. This suggests that, at least for the metagenomic and pangenomic reference collections where many references share similar *k*-mer content, better read-mapping heuristics and algorithms that improve specificity (i.e., reduce the spurious mapping of reads not arising from indexed references) without trading-off too much recall are still sorely needed. Here, it will be desirable to search for methods that can improve specificity without the need to retain reference positions or issue locate queries for all *k*-mers. We suggest that there may be several promising directions. For example, one may consider enforcing local structural consistency among matched *k*-mers to potentially reduce spurious mapping. Likewise, one may consider filtering repetitive and low-complexity *k*-mers from contributing to the final pseudoalignment result. Finally, by analogy to BLAST [4], one may consider evaluating the likelihood that a pseudoalignment result is spurious by comparing the matching rate against against some null or background expectation to account for the fact that, in very large reference databases, a very small number of (potentially correlated) *k*-mers may be insufficient evidence to consider a query as compatible with a subset of references.

## Supplementary Material

Source code: https://github.com/jermp/fulgor.

Experiments: https://github.com/jermp/fulgor-benchmarks.

## Appendix for

### A Pseudocode for the algorithms from Section 4.1

#### Algorithm 1

The Full-Intersection algorithm for a query sequence *Q*. The algorithm uses the three index components: *𝒟* (the dictionary, mapping *k*-mers to unitigs), *B* (the bit-vector mapping from unitigs to colors), and *ℒ* (the inverted index storing the compressed colors). As discussed in Section 3.1, the dictionary *𝒟* can stream through the query sequence *Q* and collect unitig ids. The inverted index ℒ, instead, returns an iterator over a color set given the color id *c* as Iterator (*c*).

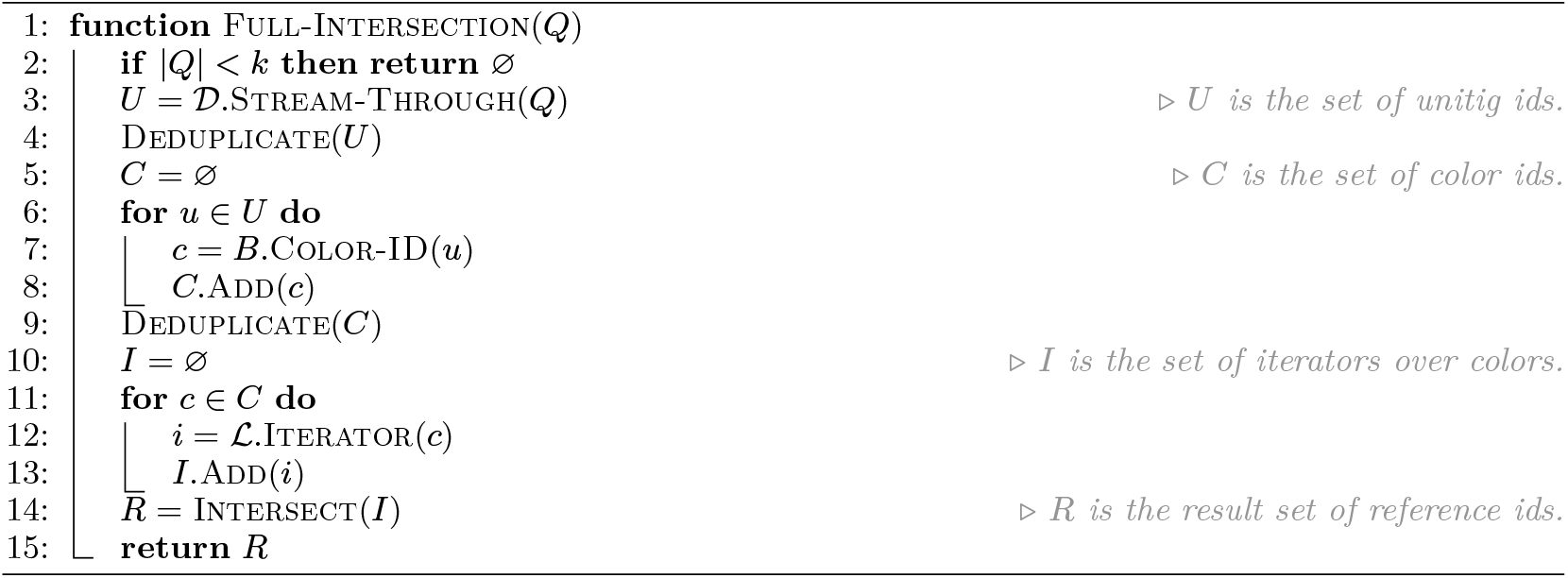

#### Algorithm 2

The Intersect algorithm for a set of iterators *I* = {*i*_1_, …, *i*_*p*_ }. An iterator object supports three primitive operations: Value (), returning the value currently pointed to by the iterator; Next (), returning the value immediately after the one currently pointed to by the iterator; Next-GEQ(*x*), returning the smallest value that is larger-than or equal-to *x*. We assume that if *i* is an iterator over color *Cj* then calling *i*.Next () for more than |*Cj*| times will return the (invalid) reference id *N* + 1.

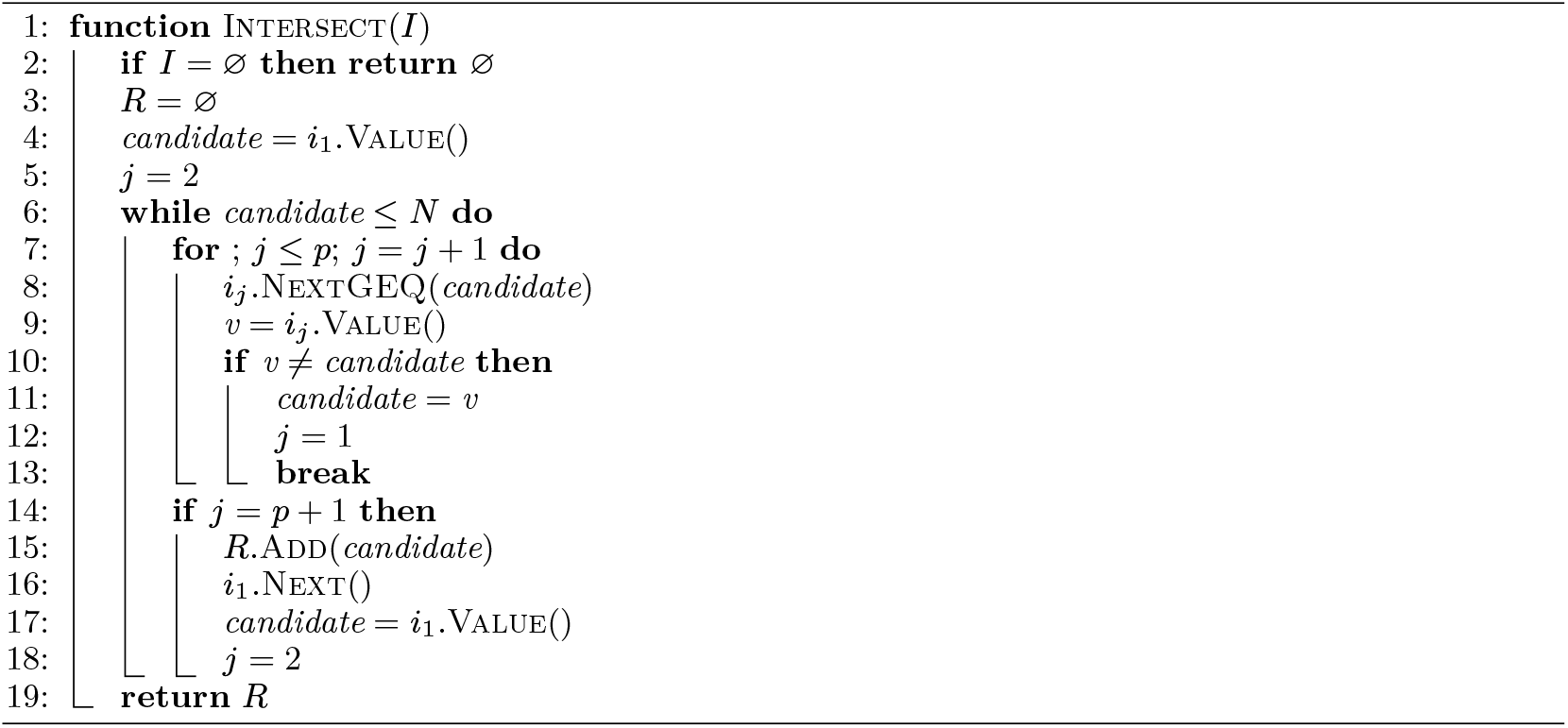

#### Algorithm 3

The Threshold-Union algorithm for a query sequence *Q*. Differently from the Full-Intersection method (Algorithm 1), here *U, C*, and *I*, are sets of pairs. The first component of a pair is a unitig id, a color id, or an iterator, respectively if the pair is in *U, C*, or *U*. The second component, read by calling the method Score () in the pseudocode, is the number of positive *k*-mers that have a given unitig id or have a given color. The score of iterator *i* is the score of the color id *c* if *i* = ℒ.Iterator (*c*). Clearly, when deduplicating the sets *U* and *C*, the scores of equal unitig or color ids must be summed.

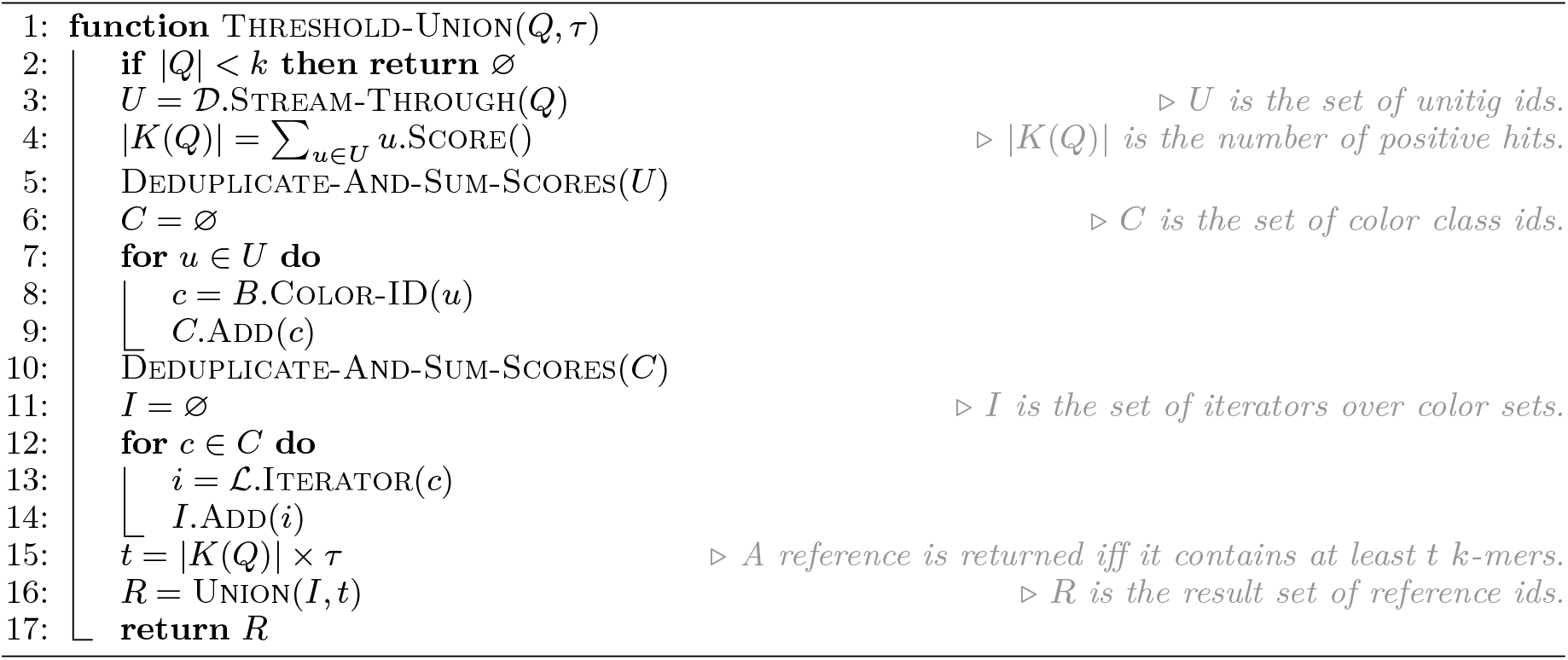

#### Algorithm 4

The Union algorithm for a set of iterators *I* = *{i*_1_, …, *i*_*p*_*}* and minimum score *t*.

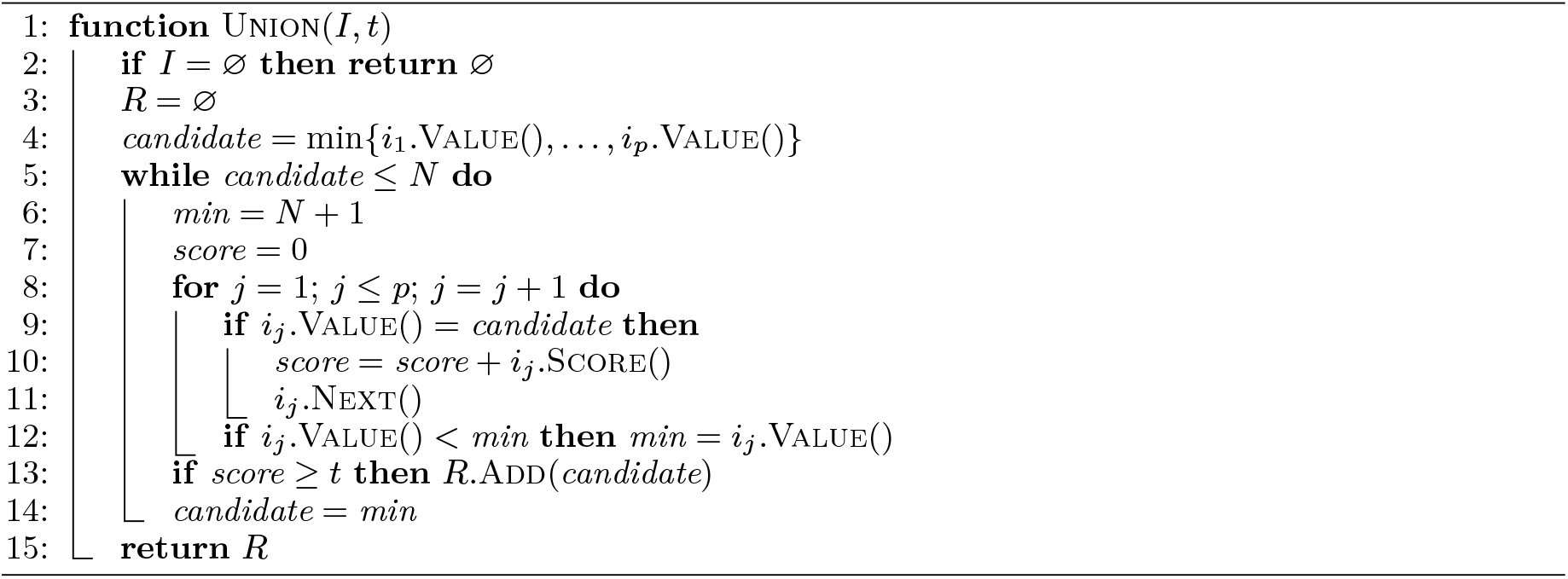

## Footnotes

1 https://github.com/jermp/fulgor/blob/main/kallisto_psa/psa.cpp

2 https://github.com/jermp/fulgor/blob/main/piscem_psa/hit_searcher.cpp

3 https://www.ebi.ac.uk/ena/browser/view/SRR896663

4 https://www.ebi.ac.uk/ena/browser/view/SRR801268

5 https://www.ebi.ac.uk/ena/browser/view/ERR321482

