## Appendix for "Fulgor: A fast and compact *k*-mer index for large-scale matching and color queries"

### A Pseudocode for the algorithms from Section 4.1

**Algorithm 1** The FULL-INTERSECTION algorithm for a query sequence $Q$. The algorithm uses the three index components: $\mathcal{D}$ (the dictionary, mapping $k$-mers to unitigs), $B$ (the bit-vector mapping from unitigs to colors), and $\mathcal{L}$ (the inverted index storing the compressed colors). As discussed in Section 3.1, the dictionary $\mathcal{D}$ can stream through the query sequence $Q$ and collect unitig ids. The inverted index $\mathcal{L}$, instead, returns an iterator over a color set given the color id $c$ as ITERATOR($c$).

```
function FULL-INTERSECTION(Q)
    if |Q| < k then return ∅
    U = D.STREAM-THROUGH(Q)                      ▷ U is the set of unitig ids.
    DEDUPLICATE(U)
    C = ∅                                        ▷ C is the set of color ids.
    for u ∈ U do
        c = B.COLOR-ID(u)
        C.ADD(c)
    DEDUPLICATE(C)
    I = ∅                                        ▷ I is the set of iterators over colors.
    for c ∈ C do
        i = L.ITERATOR(c)
        I.ADD(i)
    R = INTERSECT(I)                             ▷ R is the result set of reference ids.
    return R
```

**Algorithm 2** The INTERSECT algorithm for a set of iterators $I = \{i_1, \ldots, i_p\}$. An iterator object supports three primitive operations: VALUE(), returning the value currently pointed to by the iterator; NEXT(), returning the value immediately after the one currently pointed to by the iterator; NEXT-GEQ($x$), returning the smallest value that is larger-than or equal-to $x$. We assume that if $i$ is an iterator over color $C_j$ then calling $i$.NEXT() for more than $|C_j|$ times will return the (invalid) reference id $N+1$.

```
function INTERSECT(I)
    if I = ∅ then return ∅
    R = ∅
    candidate = i_1.VALUE()
    j = 2
    while candidate ≤ N do
        for ; j ≤ p; j = j + 1 do
            i_j.NEXTGEQ(candidate)
            v = i_j.VALUE()
            if v ≠ candidate then
                candidate = v
                j = 1
                break
        if j = p + 1 then
            R.ADD(candidate)
            i_1.NEXT()
            candidate = i_1.VALUE()
            j = 2
    return R
```

**Algorithm 3** The Threshold-Union algorithm for a query sequence $Q$. Differently from the Full-Intersection method (Algorithm 1), here $U$, $C$, and $I$, are sets of pairs. The first component of a pair is a unitig id, a color id, or an iterator, respectively if the pair is in $U$, $C$, or $U$. The second component, read by calling the method Score() in the pseudocode, is the number of positive $k$-mers that have a given unitig id or have a given color. The score of iterator $i$ is the score of the color id $c$ if $i = \mathcal{L}.\text{Iterator}(c)$. Clearly, when deduplicating the sets $U$ and $C$, the scores of equal unitig or color ids must be summed.

```
function Threshold-Union(Q, τ)
    if |Q| < k then return ∅
    U = D.Stream-Through(Q)                        ▷ U is the set of unitig ids.
    |K(Q)| = Σ_{u∈U} u.Score()                     ▷ |K(Q)| is the number of positive hits.
    Deduplicate-And-Sum-Scores(U)
    C = ∅                                          ▷ C is the set of color class ids.
    for u ∈ U do
        c = B.Color-ID(u)
        C.Add(c)
    Deduplicate-And-Sum-Scores(C)
    I = ∅                                          ▷ I is the set of iterators over color sets.
    for c ∈ C do
        i = L.Iterator(c)
        I.Add(i)
    t = |K(Q)| × τ                                 ▷ A reference is returned iff it contains at least t k-mers.
    R = Union(I, t)                                ▷ R is the result set of reference ids.
    return R
```

**Algorithm 4** The Union algorithm for a set of iterators $I = \{i_1, \ldots, i_p\}$ and minimum score $t$.

```
function Union(I, t)
    if I = ∅ then return ∅
    R = ∅
    candidate = min{i_1.Value(), ..., i_p.Value()}
    while candidate ≤ N do
        min = N + 1
        score = 0
        for j = 1; j ≤ p; j = j + 1 do
            if i_j.Value() = candidate then
                score = score + i_j.Score()
                i_j.Next()
            if i_j.Value() < min then min = i_j.Value()
        if score ≥ t then R.Add(candidate)
        candidate = min
    return R
```
